# Mechanisms of Ion Permeation in the AMPA Receptor Ion Channel

**DOI:** 10.1101/2025.06.27.662003

**Authors:** Alejandra Montaño Romero, Remy A. Yovanno, Albert Y. Lau, Edward C. Twomey

## Abstract

Excitatory synaptic transmission in the human nervous system is mediated by α-amino-3-hydroxy-5-methyl-4-isoxazolepropionic acid receptors (AMPARs), tetrameric ligand-gated ion channels localized in the excitatory post-synaptic membrane. AMPARs are activated by the binding of the neurotransmitter glutamate (Glu), which opens the ion channel and allows the influx of Na^+^ and Ca^2+^ ions into the post-synaptic neuron, initiating signal transduction. Despite many efforts, a *bona fide* ion permeation pathway of both monovalent and divalent cations in AMPARs remains elusive. From analyzing our cryo-electron microscopy (cryo-EM) map of an open calcium-permeable AMPAR (CP-AMPAR) ion channel, we identified potential sites vital to permeation of cations through the channel. To delineate mechanisms of permeation, we studied the channel with all-atom molecular dynamics (MD) simulations. Both Na^+^ and Ca^2+^ ions are coordinated by an entry site at the top of the channel prior to entering the selectivity filter. A mutation at the filter (Q607E), implicated in a neurodevelopmental disorder, makes the channel more susceptible to Zn^2+^ block but also creates a more energetically favorable environment for Na^+^ and Ca^2+^ permeation through the ion channel. These findings describe a biophysical basis for ion permeation in CP-AMPARs and how disease mutations alter the channel, which will inform therapeutic design against disease mutations in AMPARs that alter the ion channel.

## Introduction

Glutamatergic synapses are central to excitatory neurotransmission in the mammalian central nervous system (CNS)^1^. Signaling across the synapse occurs through the release of the neurotransmitter glutamate (Glu) from the pre-synaptic neuron, subsequently binding to ionotropic glutamate receptors (iGluRs) localized in the membrane of the post-synaptic neuron. There are 4 subclasses of iGluRs: a-amino-3-hydroxy-5-methyl-4-isoxazolepropionic acid (AMPA) receptors, kainate receptors, N-methyl-D-aspartate (NMDA) receptors, and delta receptors^2^. Importantly, it is the binding of Glu to AMPARs that leads to the initial depolarization of the post-synaptic neuron and downstream signaling. Given their physiological role, these receptors are critical for synaptic plasticity, memory and learning, while their dysregulation is related to developmental and intellectual delays, epilepsy, Parkinson’s disease, and additional neurological disorders^2^.

AMPARs are tetrameric multi-domain ion channels that can be made up of 4 different subunits^2,3^: GluA1 – GluA4, encoded by genes *GRIA1* – *GRIA4*. Each subunit is comprised of an amino terminal domain (ATD) that mediates receptor assembly, trafficking, and regulation, followed by the ligand binding domain (LDB) that contains the binding site for Glu, the transmembrane domain (TMD) that forms the ion channel, and the carboxy terminal domain (CTD), the intracellular domain that contributes to receptor localization and regulation. Specifically, the TMD of the AMPAR is made up three membrane-spanning helices (M1, M3, and M4), a re-entrant helix (M2) and the selectivity filter (SF). AMPARs are non-selective cation channels but are primarily permeated by Na^+^ and Ca^2+^ ions^4,5^. Moreover, Ca^2+^ selectivity is conferred by the Q/R site, a residue on the M2 helix of the GluA2 subunit that undergoes RNA editing from glutamine (Q) to arginine (R) during development; thus by adulthood the majority of GluA2 subunits are edited^6,7^. This phenomenon of GluA2 RNA editing gives rise to two main populations of AMPARs: calcium-impermeable (CI), or GluA2 containing- and calcium-permeable (CP)-AMPARs. Notably, there has been a growing appreciation of the role of CP-AMPARs in disease^8^. Interestingly, a study examining 28 patient mutations within the GluA2 subunit found a *de novo* mutation at the Q/R site, termed Q607E, mutating the codon for glutamine to one that encodes for glutamate^9^. Patients with the Q607E mutation suffer from global developmental and intellectual delays, as well as Rett-like symptoms. GluA2-Q607E AMPARs showed an increase in current amplitude, suggesting increased ion permeation, and gain of function. GluA2-Q607E AMPARs were also demonstrated to be susceptible to channel block Zn^2+^, perhaps indicating loss of function^10^.

Given their clinical relevance and clear implication in the ion permeation pathway, we decided to further investigate the molecular mechanisms of ion selectivity, permeation, and block in CP-AMPARs and how the Q607E mutation alters these processes through analyzing the permeation pathway in a cryo-EM map, and probing the pathway with all-atom molecular dynamics (MD) simulations to assess mechanisms of ion selectivity, permeability, and block of Na^+^, Ca^2+^, and Zn^2+^ ions.

## Results

### Cryo-EM reveals density in the permeation pathway

Previously, we resolved the open channel of a CP-AMPAR to 2.65 Å resolution^11^ (**Fig. 1A**). Upon further examination of the data, we observed density within the channel pore at sites that have been previously described to be critical for ion permeation^12^. Specifically, there were four main sites of interest that correspond to established binding sites. At the gate, we see density that may be coordinated by Thr617, previously reported as a Ca^2+^-specific binding site^13^. (**Fig. 1B**). Interestingly, the conditions from this structure only contain NaCl, and does not contain any significant levels of Ca^2+^.

**Figure 1.**
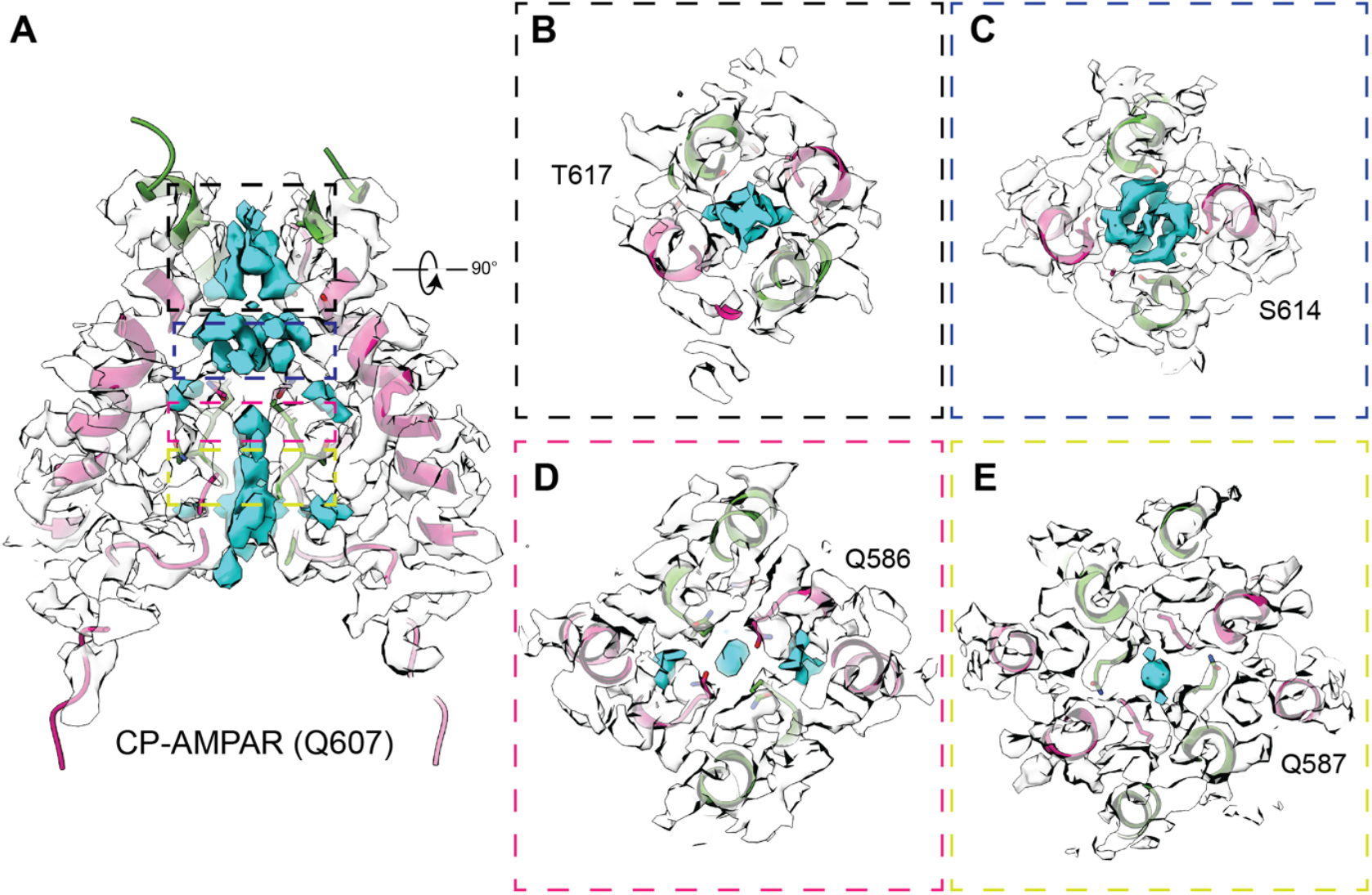
Permeation pathway density in an open CP-AMPAR ion channel. **A)** CP-AMPAR (Q607) open ion channel (M2 and M3 helices) at 2.65 Å resolution. Cryo-EM map shows density within the pore (shown in cyan) at 4 important positions, labeled by varying colored boxes. **B)** Clear density is seen at the channel gate, at Thr617. C) Density is seen below channel gate at Ser614. **C)** Density is seen at the selectivity filter, corresponding to the Gln586 side chains (also referred to as Gln607). **D)** Density seen at Gln587 backbone (also referred to as Gln608).

In the upper vestibule, we see density at Ser614, which has also been reported as an important binding site for Ca^2+^, without prior evidence in its role for Na^+^ permeation^12^. The SF is made up of the stretch of residues ^586^QQGCDI^591^ (residues 607 – 612, deviation in numbering is due to the construct used for cryo-EM^14^), where we see density within the full stretch of the SF (**Fig. 1A**). We see strong density at the central pore axis corresponding to the positions of ion permeation at the Gln586 side chain and Gln587 backbone (also referred to as Gln607 and Gln608) (**Fig. 1D-E**). These two sites have been well established to contribute to both monovalent and divalent cation permeation in CP-AMPARs^5,12,13,15^.

Thus, our data suggest a continuum of non-selective cation sites in the pore that may be vital for permeation. However, the resolution of the map does not enable direct modeling of cations or water molecules. Therefore, to uncover the thermodynamic relevance of these possible ion binding sites, we used various MD simulation methods to assess how the pathway is altered in the patient mutant AMPAR (Q607E) in the context of Na^+^, Ca^2+^, and Zn^2+^.

### Zinc acts as a pore blocker at the Q/R site

Previous literature has shown functional evidence that Zn^2+^ acts as a pore blocker in CP-AMPARs, and has higher rates of block in GluA2-Q607E patient mutant AMPARs^10^. Physiologically, Zn^2+^ is co-localized with glutamate in synaptic vesicles, making them an ion species of interest^16^. To address the molecular mechanism of Zn^2+^ channel block, we used equilibrium MD simulations involving the homomeric GluA2 open-state ion channel^14^ in either the CP (Q607) or patient mutant (Q607E) configuratio6/27/25 1:40:00 PMn. Only the ion channel helices, M2 and M3, were included and embedded within a membrane bilayer and solvated with a physiological concentration of NaCl. Given the low-level of block in CI-AMPARs^10^, we hypothesized that Zn^2+^ block was modulated by coordination via the Q/R site residues. To this end, we set up simulations with a single Zn^2+^ ion near the Q/R side chains that were run for a total of 100 ns.

For both AMPAR configurations, the Zn^2+^ ion was stabilized at the Q/R site and did not diffuse through the channel through the entirety of the simulation. Additionally, Na^+^ ions were not able to pass the Q/R site, supporting the role of Zn^2+^ as a pore blocker. For CP-AMPARs, Zn^2+^ is coordinated by the glutamine side chains in an asymmetric manner, where coordination is primarily maintained by two subunits, A and B. The distance between dimer side chains (**Fig. 2A**) shows that there is rotamer sampling occurring within the first 50 ns, primarily by the rotation of subunits C and D (**Fig. 2C**). Once the stable rotamer conformation has been established, the distance between the A/C and B/D glutamine dimers both remain ~7 – 8 Å. Interestingly, the Zn^2+^ ion translates in the X and Y planes of the pore axis during the simulation, and therefore does not maintain only one stable coordination. On the contrary, Zn^2+^ coordination in the patient mutant (Q607E) AMPAR shows a single stable orientation. Distance measurements between the glutamate side chains show that the initial rotamer is stabilized, maintaining distances of ~6 Å between B/D subunits and ~3 Å between A/C subunits (**Fig. 2B**). Side chain orientations show symmetry between A/C subunits, as they act as the primary coordinators for Zn^2+^ stabilization (**Fig. 2D**). We postulate that the increased negative charges in glutamate allow for a symmetrical orientation with increased affinity to Zn^2+^, that is not achieved by only 2 polar glutamines.

**Figure 2.**
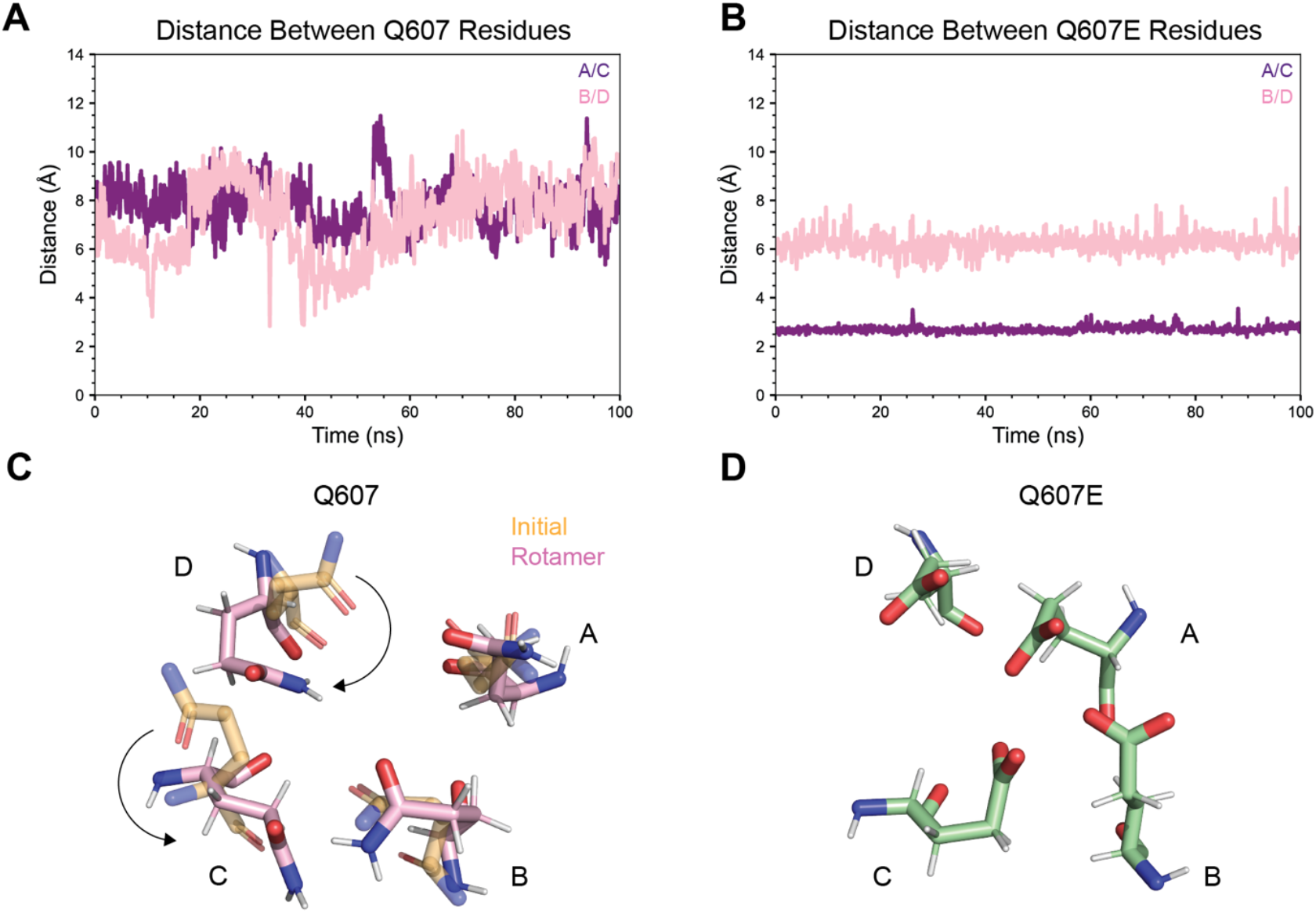
Equilibrium MD simulations show Zn^2+^ block. **A)** Distances between Q607 side chains in corresponding dimers between subunits A and C, and subunits B and D over 100 ns simulation with a single Zn^2+^ at Q/R site. Rotamer sampling of these side chains is observed over the first ~60 ns and is stabilized for the duration of the simulation. Both distances are ~8 Å in the stable confirmation. **B)** Distances between Q607E side chains between corresponding dimer subunits. Rotamers are not sampled, side chains remain stable during full duration of simulation. Distances between A and C are ~3 Å and between B and D are ~6 Å. **C)** Representation of initial glutamine side chains (yellow) to stable rotamer (pink). Zn^2+^ ion is omitted for clarity. Significant rotation is observed in subunits C and D. **D)** Representation of glutamate side chains during simulation. Zn^2+^ is omitted for clarity.

### Patient mutant Q607E AMPARs have increased ion permeation

A *de novo* patient mutation within the GluA2 ion channel mutates the glutamine at the Q/R site to a glutamate, resulting in patients that suffer from severe intellectual and developmental delays, as well as Rett-like symptoms^9^. Electrophysiology data show an increased amplitude of conductance, suggesting increased ion permeability. To determine the fundamental biophysical basis of increased ion permeability, we performed 1-dimensional (1D) umbrella sampling MD simulations to compute the free energies associated with ion permeation through the channel pore. Our umbrella sampling order parameter^17^, *χ*1, describes the longitudinal position of the ion within the central pore axis of the channel. The resulting free energy profile, or potential of mean force (PMF), gives us insights into the relative free energy of ion conduction with single-residue resolution. The umbrella sampling windows cover a 46 Å range of the ion channel and are positioned at 0.5 Å increments (**Fig. 3D**). This approach was used with Na^+^, Ca^2+^, and Zn^2+^ ions separately (**Fig. 3**).

**Figure 3.**
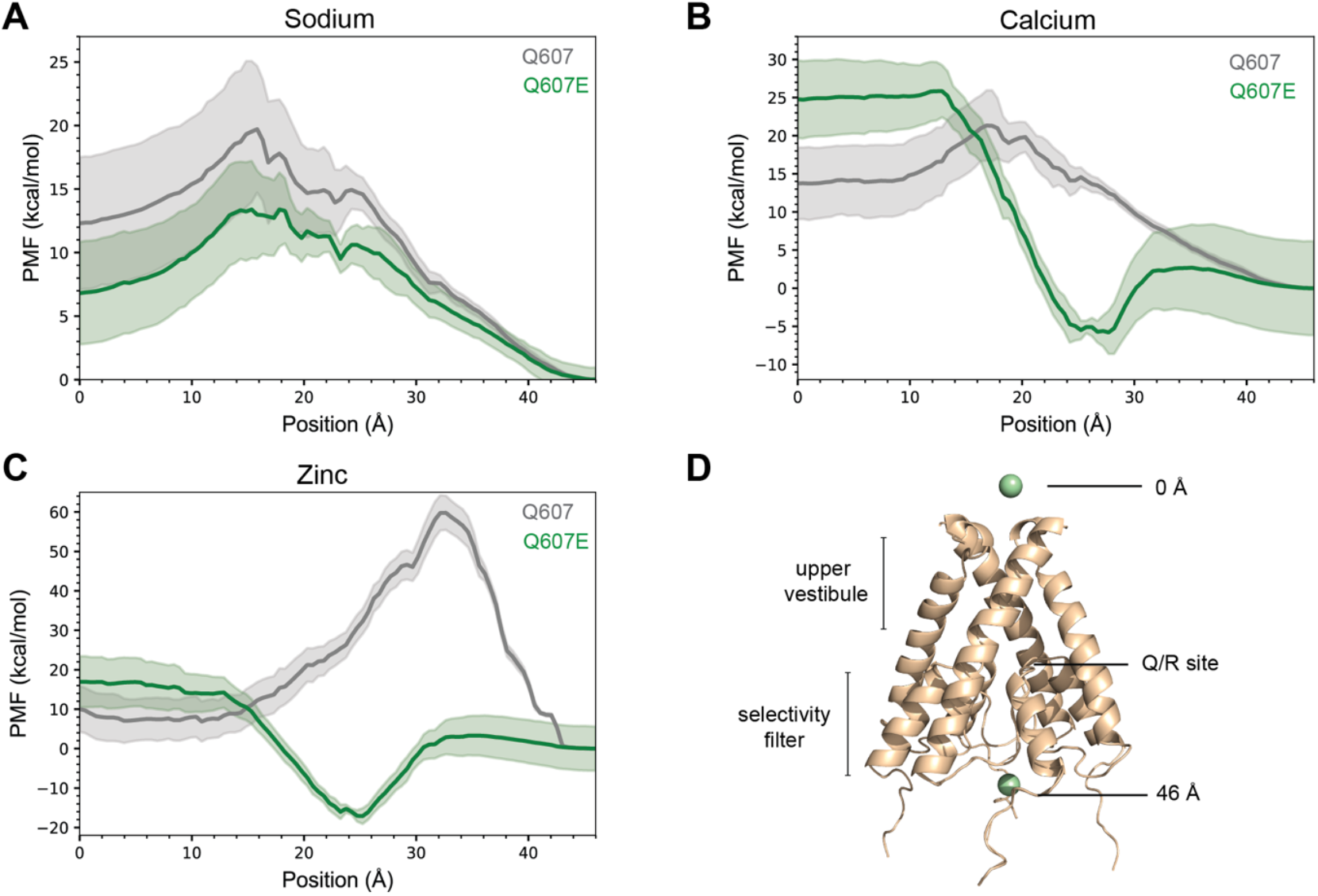
1D umbrella sampling of Na^+^, Ca^2+^, and Zn^2+^. **A)** PMF of Na^+^ in CP-AMPARs (Q607) and patient mutant AMPARs (Q607E). In the Q607 channel, the ion encounters a free energy barrier of 7.4 kcal mol^−1^ to enter the channel and is stabilized at 2 local minima corresponding to site G (position ~17 Å) and the Q/R site (position ~23 – 24 Å) along its permeation pathway. In the Q607E channel, there is a smaller free energy barrier to enter the channel of 6.2 kcal mol^−1^ and a single local minimum at the Q/R/E site during permeation. **B)** PMF of Ca^2+^. In Q607, the ion encounters a free energy barrier at *χ* ≈ 18 Å and is stabilized at the position corresponding to site G via a hydrogen bonding network during permeation. In Q607E, the PMF suggests that the ion favorably enters the pore, descends to the Q/R/E site and is displaced by a second Ca^2+^ ion to finish permeation through the channel. **C)** PMF for Zn^2+^. In the Q607 channel, Zn^2+^ encounters a significant barrier to enter the pore, suggesting that entrance is voltage dependent. In the Q607E channel, Zn^2+^ favorably enters the pore and remains coordinated at the Q/R/E site without permeating through. Additionally, the trend suggests there is less voltage-dependence to enter the pore. **D)** Representation of AMPAR ion channel with only M2 and M3 helices. *χ* = 0 Å and 46 Å, indicated by the ion at the central pore axis.

For Na^+^ with CP-AMPARs (Q607), there is a free energy barrier of 7.4 kcal mol^−1^, from 0 to ~16 Å, for the Na^+^ to enter the pore. Once Na^+^ overcomes the barrier to enter the channel, the decreasing PMF suggests favorable permeation (**Fig. 3A**). Interestingly, we see 2 local minima that correspond to the gate residue Thr617 (*χ* ≈17 Å) and the Q/R site (position *χ* ≈ 23 – 24 Å). Local minima indicate metastable sites during permeation, suggesting that the Na^+^ ion is coordinated by the residues at the gate and the Q/R site. Furthermore, in the patient mutant AMPAR (Q607E), the Na^+^ must also surmount a free energy barrier, albeit a smaller one, of 6.2 kcal mol^−1^. The overall remaining trend of the free energy landscape supports permeation in the mutant channel, though with only a significant local minimum corresponding to the Q/R/E site. Taken together, the data suggest that when compared to Q607, the Na^+^ undergoes less of an energy barrier to enter the Q607E pore, with a ΔΔ*G* of –1.2 kcal mol^−1^. Additionally, the permeation pathway of Na^+^ differs slightly in the Q607E pore, where the ion primarily is stabilized at the Q/R/E site, and not Thr617, during permeation.

Umbrella sampling simulations with Ca^2+^ show two distinct free energy landscapes, both indicating favorable permeation but by different mechanisms (**Fig. 3B**). In CP-AMPARs (Q607), the trend is reminiscent of that seen in Na^+^, where Ca^2+^ surmounts a free energy barrier to enter the pore, from *χ* ≈ 0 to 16 Å. The Ca^2+^ ion then has two free energy minima, corresponding to Thr617 and the Q/R site, before the continued free energy descent along the remaining selectivity filter. The coordination at Thr617 is mediated by a hydrogen bonding network with waters in the pore, therefore giving us a local free energy minimum at *χ* ≈ 18 Å. The importance of Thr617 and its interactions within a hydrogen bonding network has been described previously for the Ca^2+^ permeation pathway, in both molecular dynamics simulation and cryo-EM studies^12,13^. Strikingly, we see a drastic change in the free energy landscape in patient mutant (Q607E) AMPARs. Under this configuration, Ca^2+^ does not need to overcome any energy barrier to enter the pore, and rapidly undergoes a free energy descent from *χ* ≈ 13 Å to the Q/R/E site at *χ* ≈ 23 – 24 Å. In these simulations, we observe the side chains of Thr617 in subunits A and C directly coordinate Ca^2+^ at *χ* ≈ 13 Å before the ion moves to the Q/R/E site. The landscape has a well that encompasses the Q607E side chain and the Q608 (also referred to as Q587) backbone, suggesting that the ion can be stabilized at these positions, as previously reported^12^. Finally, the Ca^2+^ must cross a small energy barrier to finish translocating through the pore. Given this overall pattern, we propose that there is a significant increase of Ca^2+^ permeation in the Q607E AMPARs via a local concentration effect. The electrostatic charge within the pore attracts Ca^2+^ ions, allowing them to easily enter the pore and move to the Q/R/E site, which is translocated by the entrance of a second Ca^2+^ ion, following a knock-on mechanism.

### Patient mutant Q607E reduces voltage-dependence of zinc channel block

Our equilibrium MD simulations gave insights into the mechanism of Zn^2+^-mediated channel block. To further investigate, we performed 1D umbrella sampling with Zn^2+^. Given our results of rotamer sampling in Q607 AMPARs, we used the stable rotamers induced by ~60 ns of simulation as the conformation used for the umbrella sampling simulations (**Fig. 2C**). As expected, the PMF indicates Zn^2+^ block, rather than permeation, but with distinct mechanisms for each AMPAR configuration (**Fig. 3C**). In CP-AMPARs (Q607), there is a significant free energy barrier for entrance and translocation through the pore, with an energy maximum at *χ* ≈ 32 Å. The data suggest that it is not favorable for the Zn^2+^ ion to enter the channel. A caveat of these simulations is that there is no applied electric potential, or a force that recapitulates a biological membrane voltage potential, suggesting that Zn^2+^ entrance and block of Q607 AMPARs are voltage dependent. On the contrary, in patient mutant (Q607E) AMPARs, Zn^2+^ does not have an energy barrier to enter the channel. Instead, it easily makes it to the Q/R/E site, represented by the free energy minimum, and is stably coordinated at that site. This model is supported by the nearly equal free energy barriers from the Q/R/E site to either the top (0 Å) or the bottom (46 Å) of the channel. Additionally, the data suggest that Q607E renders Zn^2+^-mediated block to be less voltage-dependent.

### Mechanism of ion permeation

Given the results from re-analyzing our cryo-EM map and MD simulations, we propose a mechanism of both increased ion permeation and block in the Q607E AMPARs when compared to CP-AMPARs, where the changed electrostatics alter the energetics of the pathway (**Fig. 4**). Our data suggest that Q607E mutant receptors undergo increased Na^+^ permeation by decreasing the energy barrier for the ion to enter the channel and is stabilized at the Q/R/E site before permeating through the channel. Due to the drastic difference observed in the PMF of Ca^2+^, we infer increased Ca^2+^ mediated conductance by a local concentration effect of the four glutamates at the Q/R/E site. Finally, our data suggest that there are increased levels of Zn^2+^ mediated block in Q607E receptors through a more stable coordination of Zn^2+^ at the Q/R/E site and a decreased voltage-dependance for entrance into the channel.

**Figure 4.**
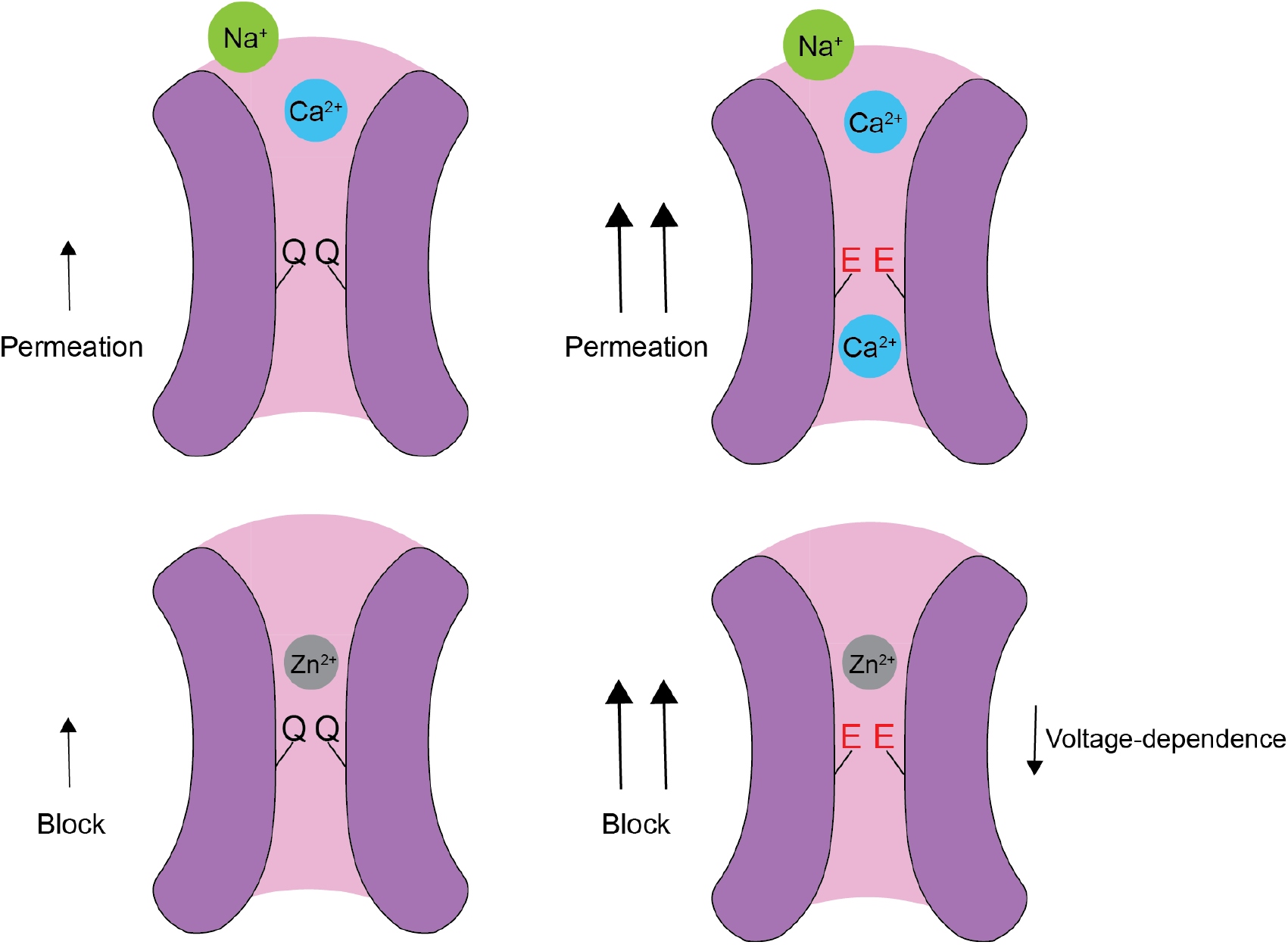
Mutant Q607E effects on ion permeation and block. Model on patient mutant (Q607E) effects on ion permeability and block. Q607E increases Na^+^ and Ca^2+^ permeability through electrostatic changes, while also increasing Zn^2+^ block by increased affinity and lower voltage-dependence for Zn^2+^ entry.

## Discussion

Regulation of ion permeation, particularly that of Ca^2+^, is vital for proper neuronal signaling and function. This is exemplified by the developmental role of RNA editing of the GluA2 subunit Q/R site, that provides a modification to the levels of Ca^2+^ permeability of AMPARs in adulthood^6,7^. Deviation from this balance by the increase of CP-AMPARs populations, whether through subunit expression levels or ion channel mutations, leads to a plethora of aberrant physiological states. A patient mutation at this Q/R/E site in GluA2, termed Q607E, leads to global developmental and intellectual delays along with a Rett-like symptoms^9^. In this work, we show through previous cryo-EM and MD simulations that the electrostatic charge difference between calcium-permeable (Q607) and patient mutant (Q607E) alters the thermodynamics of the AMPAR ion permeation pathway.

Mutations within the GluA2 subunit have global effects as the GluA2 mRNA is expressed in all brain regions and is accompanied by another AMPA receptor subunit transcript to form heteromers – therefore, GluA2-containing heterotetramers are the most common AMPARs in the CNS^2,18^. The GluA2 subunit determines numerous functional properties of AMPARs and its expression influences circuit function^2^. Together, this highlights the physiological gravity of a patient mutation within the GluA2 subunit and is further emphasized by one that affects the ion permeation pathway.

Our work shows how the ion permeation pathway is altered for both monovalent and divalent cations, in the context of Na^+^, Ca^2+^, and Zn^2+^. In CP-AMARs, Na^+^ encounters two main binding sites during permeation: 1) at the gate at Thr617 and 2) the Q/R/E site residue side chain. Interestingly, the binding site at Thr617 has only previously been observed for Ca^2+^, but our data suggest that this site might be a universal cation binding site^13^. In the mutant Q607E-AMPAR, there is increased Na^+^ permeation via a reduction of the ion’s free energy barrier to enter the pore and is primarily stabilized by the Q/R/E side chain. With Ca^2+^, the thermodynamics of ion permeation is dramatically altered by the change in electrostatics within the inner portion of the channel. With two glutamates present, we postulate that an increased local concentration of Ca^2+^ develops, like what has been seen and established by the DRPEER motif in NMDARs^19^. Though the DRPEER motif in NMDARs is in the extracellular vestibule and the Q/R/E site is below the channel gate, our data suggest that a similar effect occurs in Q607E-AMPARs that facilitates the translocation of Ca^2+^ to the Q/R/E site^19^. Increased Ca^2+^ permeation is known to trigger signaling events in the post-synaptic neuron that are accompanied by changes in synaptic efficacy and neuronal morphology^2^. Finally, increased Zn^2+^ block is seen in Q607E-AMPARs with less voltage-dependence for entrance of Zn^2+^ into the channel. Overall, this model shows how the Q607E-AMPAR is a combination of gain-of-function permeation for Na^+^ and Ca^2+^, and loss-of-function permeation for Zn^2+^, causing a severe imbalance in AMPAR function.

In conclusion, we outline clear ion permeation pathways for both monovalent and divalent cations and how they are altered between CP-AMPARs and Q607E-AMPARs with single-residue resolution. Precision structural biology, such as that presented in this work, with patient mutations inform the design of specific therapeutics for fine-tuning AMPAR function.

## Methods

### Equilibrium molecular dynamics

A construct of the tetrameric M2-M3 helices (residues 565 – 624) of the GluA2-Q607 was built from the open-state cryo-EM structure^20^ (PDB ID: 5WEO). PyMOL was used for single-residue mutagenesis at Q607 (equivalent to residue 586) for the GluA2-Q607E construct. The constructs were embedded in a POPC membrane using CHARMM-GUI Membrane Builder^21,22^. All systems were solvated in a 100 Å x 100 Å x 105 Å orthorhombic water box with 150 mM NaCl using CHARMM. The system was electrically neutral. All simulations described in this work were performed using the CHARMM36 forcefield^23,24^ with explicit solvent at 300 K. The all-atom potential-energy function PARAM27 for proteins^25,26^ and the TIP3P potential-energy function for water^27^ were used. Both the N- and C-termini were capped with ACE and CT3 terminal group patching, respectively. Equilibration was carried out in the NVT ensemble with restraints applied to the backbone and side chain atoms and were gradually released over the course of equilibration. Production simulations were carried out in the NPT ensemble at 1 atm and 300 K. Long-range electrostatic interactions were computed using the particle mesh Ewald algorithm^28^. NAMD was used for all simulations^29^.

Simulations with a single zinc ion within the pore were set up as previously described except for using PyMOL to add in the zinc ion at a specified location during construct generation. The simulations were electrically neutral.

Simulations with 50 mM extracellular ZnCl2 were established as previously described except for solvating with 50 mM ZnCl2 in place of 150 mM NaCl. The simulations were electrically neutral.

### 1D umbrella sampling

The conformational free energy landscapes, or potentials of mean force (PMFs), of ion permeation were computed using 1-dimensional (1D) umbrella sampling simulations^17^. A single order parameter (*χ*) described the position of an ion along the channel. Coordinates for the umbrella sampling windows were generated by translocating the ion along the desired Z-range in PyMOL. Windows were positioned in 0.5 Å increments over a total of 46 Å, resulting in a total of 94 windows. Each window was solvated in a 100 Å x 100 Å x 105 Å orthorhombic water box with 150 mM NaCl using CHARMM, creating an electrically neutral system. Simulations used the CHARMM36 forcefield with explicit solvent at 300 K. The all-atom potential-energy function PARAM27 for proteins and the TIP3P potential-energy function for water were used. Both the N- and C-termini were capped with ACE and CT3 terminal group patching, respectively. Equilibration was carried out in the NVT ensemble with restraints applied to the backbone and side chain atoms and were gradually released over the course of equilibration. Production simulations were carried out in the NPT ensemble at 1 atm and 300 K. Long-range electrostatic interactions were computed using the particle mesh Ewald algorithm.

Umbrella sampling simulations were performed by applying a harmonic restraining potential with a force constant of 20 kcal mol^−1^ Å^−2^ to *χ* for each of the 94 umbrella windows. To maintain the open state conformation, a 100 kcal mol^−1^ Å^−2^ harmonic center-of-mass restraint was added to the N, CA, and C atoms of residues 565 and 624^30^. Each PMF comprises 94 windows totaling 470 ns of simulation time. The biased sampling was mathematically unbiased and combined using the weighted histogram analysis method (WHAM)^31^. The error was computed by calculating standard deviations by block averaging with ten blocks of sampling for each window^32^.

## Acknowledgements

We thank members of the Twomey and Lau labs for helpful discussions during the development of this work. Computational resources were provided by the Maryland Advanced Research Computing Center (MARCC) and Advanced Research Computing at Hopkins (ARCH) at Johns Hopkins University. A.Y.L. is supported by National Institutes of Health (NIH) grant R01GM094495. E.C.T. is supported by NIH grant R35GM154904 and the Searle Scholars Program (Kinship Foundation #22098168).

## Competing Interests

The authors declare no competing interests.

## Notes

### Competing Interest Statement

The authors have declared no competing interest.

